# A scalable recombinant pipeline for disulphide-stapled peptides

**DOI:** 10.64898/2026.09.29.755317

**Authors:** Yensi Flores Bueso, Gizem Gökçe-Alpkılıç, Jihun Jeung, Stephen Rettie, Xinting Li, Mark Tangney, David Baker, Gaurav Bhardwaj

**Author notes:** Equal authorship (Yensi Flores Bueso, Gizem Gökçe-Alpkılıç). Correspondence: Yensi Flores Bueso.

## Abstract

Constrained macrocyclic peptides can engage protein surfaces that resist both small molecules and biologics but producing them at library scale still depends on chemical synthesis and macrocyclisation workflows that are slow, specialised and hard to parallelise. We describe a recombinant pipeline for disulphide-stapled cyclic peptides, compatible with 96 well plates, that lowers this barrier. Peptide-encoding sequences are introduced by extension PCR and in vivo assembly (IVA) cloning at about $2 per variant, expressed as small ubiquitin-like modifier (SUMO) fusions secreted to the oxidising periplasm, where the intramolecular disulphide forms spontaneously, and purified by immobilised metal-affinity chromatography (IMAC) with analytical size-exclusion chromatography (SEC). The route from a list of designs to characterised material takes about one week, much of it unattended. Applied to a 60-design library across nine protein targets, the pipeline gave purified products for all 60 designs, at a median total soluble yield of approximately 135 micrograms per 6 mL culture (range 9 to 261) that varied by target. Ellman’s assay across 38 designs showed disulphide-formation failure to be sequence-driven and uncorrelated with yield, detecting a failure mode invisible to chromatography. Surface plasmon resonance confirmed target binding and showed that the recombinant SUMO fusion constructs retained affinities comparable to their corresponding peptide-only forms. Complementary to general-purpose recombinant platforms, the pipeline enables higher-throughput disulphide macrocycle screening and is amenable to partially or fully automated workflows.

## INTRODUCTION

Many therapeutically significant protein targets feature recognition sites that are broad and relatively topographically featureless, making them difficult to target with conventional small molecules^1^. At the other end of the spectrum, protein therapeutics can achieve high specificity, but their size and physicochemical properties often constrain delivery and limit access to intracellular space^2^. Constrained peptides, particularly macrocycles, lie between these two modalities: they are compact and they can now be designed against targets to present a defined binding surface that engages the extended contact regions typical of protein–protein interactions with precision^3–6^. Constrained macrocycles could therefore reach target classes that remain challenging for both small molecules and biologics^7^.

Despite this potential, producing macrocyclic peptide libraries at scale remains a practical bottleneck. Chemical synthesis and macrocyclisation workflows are time-intensive, specialised, and difficult to parallelise without dedicated, high-end infrastructure, which restricts routine use to laboratories with substantial technical capacity and equipment access^7, 8^. In addition, throughput is constrained by iterative synthesis, cyclisation optimisation, and complex purification, which together increase the cost and turnaround time. These workflows also rely on substantial volumes of organic solvents and other hazardous reagents, increasing the environmental footprint and the waste-handling burden^9^. As a result, large-scale generation and validation of macrocyclic libraries can be slow and cost-prohibitive, limiting broad access to the technology and slowing iterative design–test cycles. Lowering these practical barriers would make macrocycle discovery campaigns feasible in a wider range of academic settings. There is therefore a need for a reproducible, mid- or high-throughput pipeline that enables parallel production, purification, and quality control of macrocyclic variants at lower cost and without requiring highly specialised equipment.

Genetically encoded disulphide macrocycles have several practical advantages for iterative design and screening. The macrocyclisation chemistry is encoded directly in the sequence and can occur spontaneously within the oxidising environment of the bacterial periplasm, bypassing the need for exogenous catalysts or post-synthetic modification^10^. Recombinant production lowers the cost and entry barrier and supports parallelisation, enabling rapid iteration across variant libraries and simplifying scale-up. In addition, solubility and affinity tags streamline purification and support automation (e.g., immobilised metal-affinity chromatography (IMAC) workflows) and increase the effective molecular mass of the analyte, which can improve signal-to-noise in label-free binding measurements (e.g., surface plasmon resonance SPR) and reduce the amount of material required for reliable kinetic screening^11^. Where needed, tags can be cleaved. Finally, because these peptides are genetically encoded and recombinantly produced, they provide a viable architecture for live biological therapeutics (LBTs)^12^. This opens a path for *in situ* delivery within host microbiomes, such as the targeted secretion of antimicrobial macrocycles, which could be advantageous for ecological or clinical interventions where traditional administration is unfeasible^13^.

Here, we establish a reproducible 96-well-compatible pipeline combining IVA cloning^14^, periplasmic expression, tag-assisted purification, and rapid analytical QC (size-exclusion chromatography (SEC) and Ellman’s assay, with confirmatory mass spectrometry (MS) and binding measurements where required) to enable mid-throughput production and evaluation of disulphide-stapled cyclic peptides.

## RESULTS

### 1. Overview of the mid-throughput pipeline and performance targets

We designed an end-to-end workflow integrating automated primer design, *in vivo* assembly (IVA) cloning, periplasmic expression with a small ubiquitin-like modifier (SUMO) solubility/capture fusion, IMAC, and analytical size-exclusion chromatography for parallel processing of up to 96 designs (Fig. 1A). The workflow delivers purified, characterised peptide-fusion product in less than a week after ordering primers (Fig. 1B). Much of this week is passive: standard <60-nucleotide primer pairs are synthesised and delivered in ~2–3 days (against ~8 days for ~300-bp gene fragments used conventionally) after which cloning, expression and purification can proceed within 3-4 days with limited hands-on time. By replacing synthetic gene fragments (eBlocks at list price ~$0.07 per base for ~300-bp constructs) with standard < 60-nucleotide oligonucleotide primer pairs that introduce the peptide-encoding sequence by extension PCR, the per-variant DNA cost is reduced to approximately $2 per design (Fig. 1C), which is about an order of magnitude lower than gene-fragment input, while remaining compatible with low-volume parallel cloning in 96-well format.

**Figure 1.**
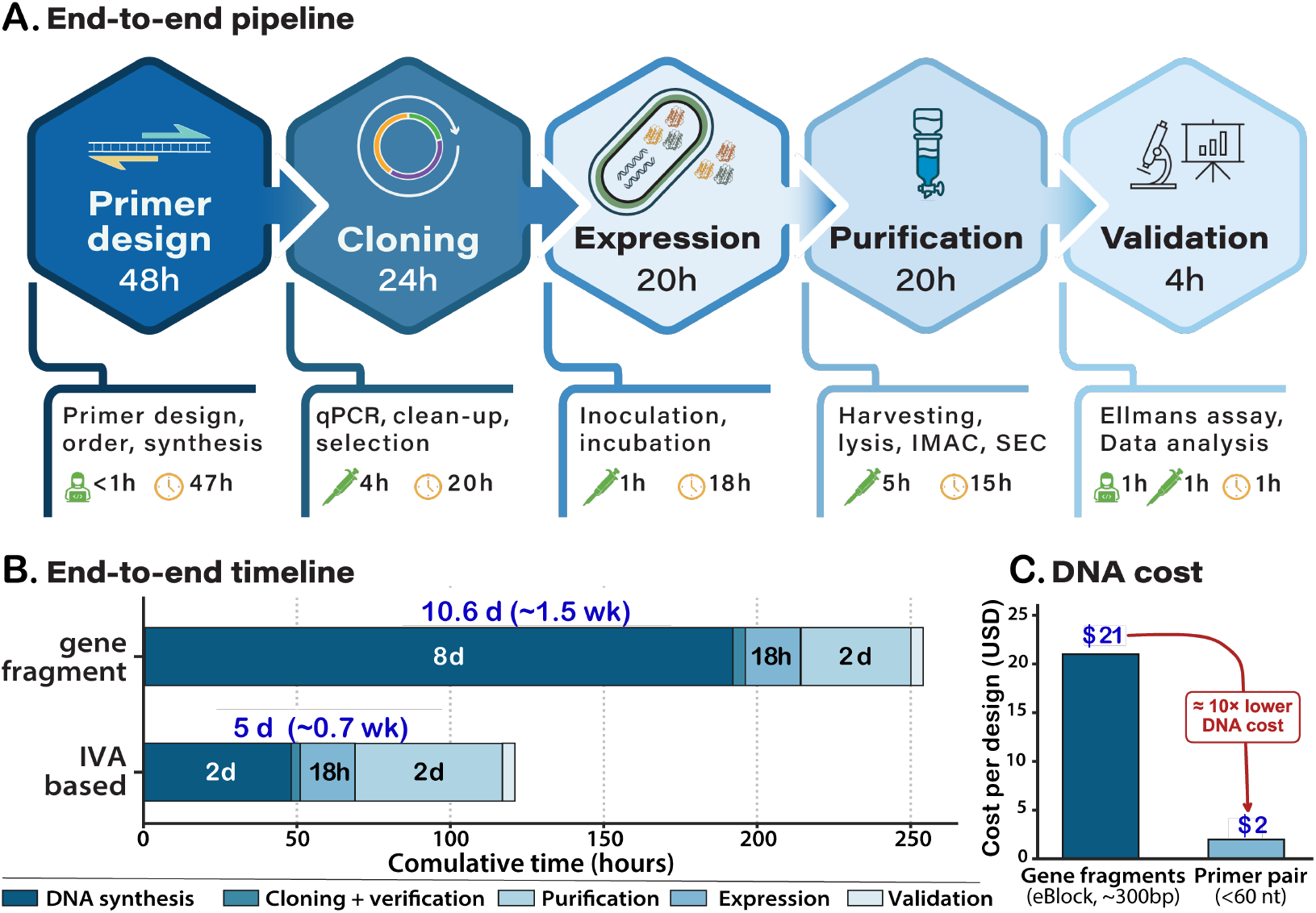
A 96-well recombinant pipeline for disulphide-stapled cyclic peptides: workflow, timeline and cost. **(A) End-to-end workflow**: automated primer design, *in vivo* assembly (IVA) cloning, periplasmic expression of a Small Ubiquitin-like Modifier (SUMO)–peptide fusion, immobilised metal-affinity chromatography (IMAC), and analytical size-exclusion chromatography (SEC), processing up to 96 designs in parallel. **(B) Time from a list of designed sequences to characterised**, purified material (~1 week, ~3-4 working days) versus conventional gene-fragment routes (~1.5 weeks). The DNA-synthesis block is vendor turnaround; passive waiting for oligo (~2–3 d) or gene-fragment (~8 d). **(C)** Per-variant DNA cost: standard oligonucleotide primer pairs (~$2 per design) versus synthetic gene fragments (~$0.07 per base for ~300-bp constructs).

#### The pipeline is modality-specific

General-purpose recombinant pipelines such as SAPP/DMX^15^ target arbitrary soluble-protein designs at hundreds of designs per day and ~$5 per construct via DNA oligo demultiplexing. Instead, our workflow integrates the steps that disulphide-stapled cyclic peptides require: a SUMO fusion that both enhances expression of cysteine-rich short sequences and provides the IMAC tag; periplasmic secretion to reach the oxidising environment needed for spontaneous intramolecular disulphide formation; and an Ellman’s-based read-out for staple formation alongside SEC for yield and dispersity. Together these give a compact, lower-cost and more sustainable pipeline that produces and quality-controls disulphide-stapled cyclic peptides on standard molecular-biology equipment, without specialist and solvent-heavy peptide-synthesis infrastructure^9^, lowering the barrier for academic labs to run iterative design–test campaigns.

In the sections below we describe the development and optimisation of each step: primer design and cloning (2), expression and lysis (3). We later apply the full workflow to a library of 60 designs targeting nine distinct protein targets and present the yield analysis (4), and disulphide quality control (5). We finally confirm that pipeline material retains binding affinity comparable to the corresponding chemically synthesised peptide of the same sequence (6).

### 2. Automated Design and High-Throughput Cloning

#### Automated Design

To support routine 96-well cloning, we developed a pipeline that automates primer design from input amino-acid sequences (see methods 1.2). For each design, the pipeline reverse-translates the peptide sequence with codon optimisation for *E. coli*, applies sequence-level constraints (GC content ~60%, avoidance of homopolymers and predicted mRNA structures around the ribosome-binding site, removal of internal restriction sites), and assembles the IVA primer pair from three concatenated regions: a fixed 3′ adapter tail (Tm 63 °C) for template annealing, a central peptide-encoding extension, and a 5′ homology block of variable length (14–20 nt) tuned to a Tm window of 48–56 °C to support efficient RecA-independent recombination. The pipeline outputs primer-plate maps, theoretical molecular weights and lengths for each downstream peptide–fusion product, and order-ready CSVs in the format required by commercial primer suppliers. We used this pipeline to design and order all primers reported in this work (see extended data).

#### Cloning

We optimised the IVA amplification by qPCR to set a reliable threshold before transformation and reduce the amount of reagent required. PCR products were monitored in real time using EvaGreen-supplemented Q5 Master Mix on a CFX96 instrument; 10 μL reactions reaching exponential amplification before cycle 15 with >5,000 RFU were taken forward to transformation, as those that reach exponential amplification at later cycles yielded no transformants (Fig. 2B). Across the libraries reported here, amplification efficiency was peptide-dependent rather than primer-design-dependent, with ~95% (91/96) of reactions meeting the threshold within 15 cycles. Reactions below the threshold were excluded.

**Figure 2.**
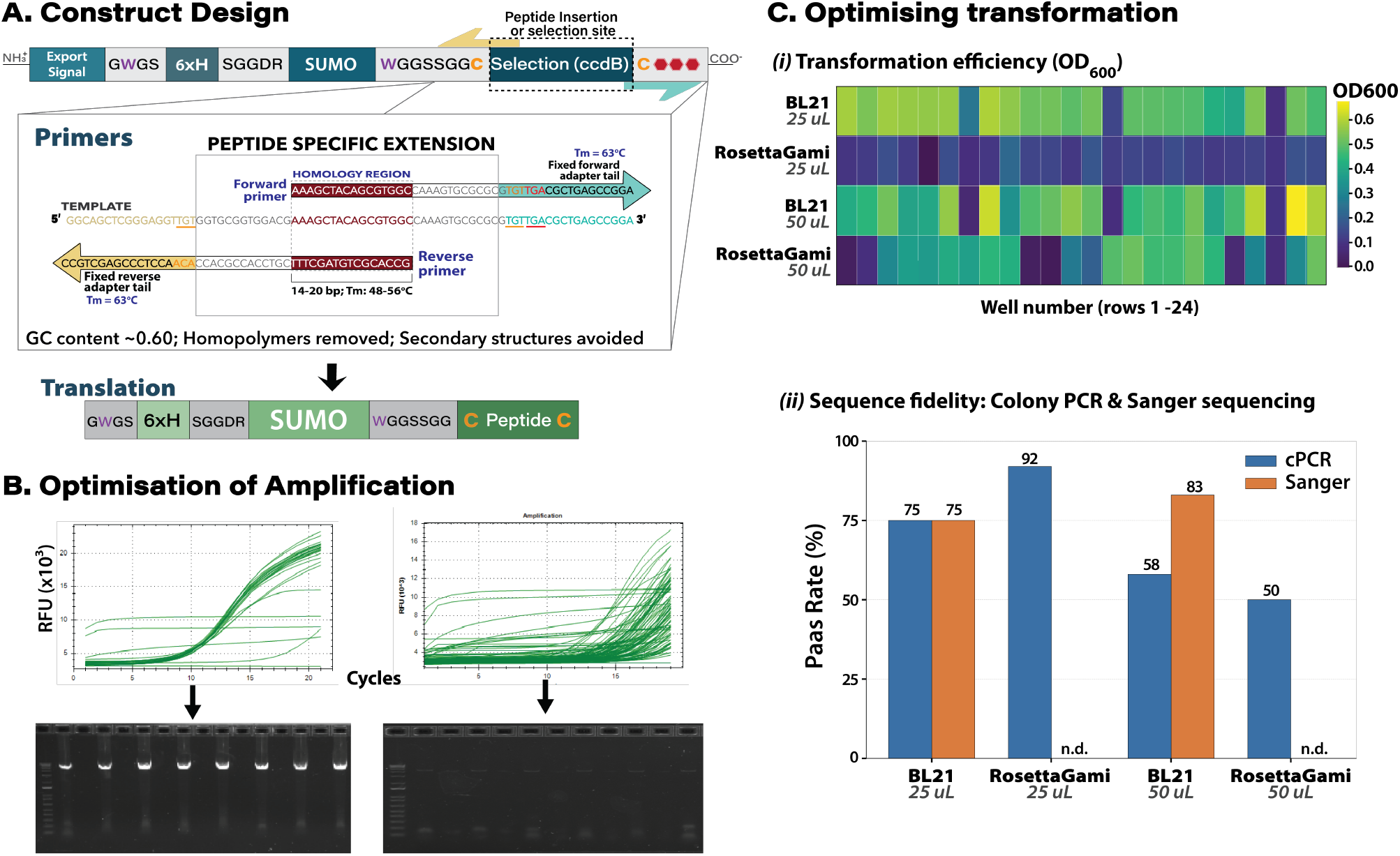
Construct design and cloning efficiency. **(A) Illustrative expression construct** (His6–SUMO– linker–peptide in a pET-24b(+) backbone), with the ccdB negative-selection cassette excised during IVA cloning. **Primer architecture:** a fixed 3′ adapter tail (yellow: reverse, cyan: forward), a peptide-encoding extension (dashed box), with a 5′ homology region (red) and a representative peptide sequence in grey. **(B) Representative real-time PCR (qPCR) amplification traces** and the agarose electrophoresis gels of a colony PCR using T7 promoter primers (~1000 bp) to verify cloning; ***(left)*** reactions reaching exponential phase before cycle 15 and above 5,000 relative fluorescence units (RFU), were taken forward to transformation; ***(right)*** reactions reaching exponential phase later than ~cycle 17 rarely produced transformants and were flagged and excluded. **(C) Transformation optimisation**: ***(i)*** efficiency (OD600) across four conditions × 24 wells: two strains (BL21 *DE3*, Rosetta-Gami), two cell volumes (50 and 25 μl) and two PCR-product volumes (5 and 2 μl); the highest transformation efficiency was observed for BL21(DE3). ***(ii)*** Sequence fidelity: we assessed cloning quality via colony-PCR (cPCR) and subsequent sanger-sequencing of the amplicon; the standard adopted was 25 μl BL21(DE3) with 2 μl of DpnI-treated PCR product (“n.d.”, not determined).

#### Transformation

We screened a 96-reaction (four-condition) matrix of two competent-cell strains (E. coli BL21(DE3) and Rosetta-Gami) at two transformation volumes (25 and 50 μl), 24 wells per condition (Fig. 2C). After overnight outgrowth in selective LB-kanamycin we measured culture density (OD600, Fig. 2C-i) and confirmed insert presence by colony PCR (cPCR). BL21*(DE3)* transformed far more consistently than Rosetta-Gami, which gave low culture density across most wells at both volumes. Pass rates were calculated over the wells that transformed; they therefore report sequence fidelity conditional on successful transformation. On that basis the cPCR pass rate was highest for Rosetta-Gami at 25 μl (92%), followed by BL21 at 25 μl (75%) and 50 μl (58%), with Rosetta-Gami 50 μl lowest (50%; Fig. 2C-ii). Sanger sequencing of cPCR-positive colonies confirmed insert identity in 75% (BL21, 25 μl) and 83% (BL21, 50 μl) of sampled wells (not determined for Rosetta-Gami). Because Rosetta-Gami rarely produced transformants despite high per-colony fidelity, we adopted BL21(DE3) at 25 μl as the standard with 2 μl DpnI-treated PCR product per transformation. The cell-aliquot volume was subsequently reduced to a working minimum of 15 μl (10, 15, 20 and 25 μl were tested, and 15 μl was the lowest that transformed reliably) and used as the standard for all subsequent libraries. Under this condition the cloning step from PCR setup to selected overnight culture takes about 4 hours of hands-on time per 96-well plate.

### 3. Optimisation of Secretion and Periplasmic Extraction

The *E. coli* cytoplasm has a reducing environment, which is incompatible with stable intramolecular disulphide formation, leading to toxic free-cysteine-rich sequences accumulating in the cytosol^10^. In early attempts at cytoplasmic expression of the SUMO-stapled peptide constructs in BL21*(DE3)* we observed little soluble product across multiple conditions, as judged by SEC traces and SDS-PAGE of clarified lysates. We therefore aimed for expression to the periplasm, where the oxidising environment supports spontaneous disulphide-bond formation through endogenous periplasmic thiol-disulphide oxidoreductase (Dsb) activity.

To find a secretion signal that would translocate the SUMO-peptide fusion across the inner membrane, we screened seven N-terminal secretion signals: DsbA, MalE (maltose-binding protein), OmpA (outer-membrane protein A), PhoA (alkaline phosphatase) and PelB (pectate lyase B), together with the vesicle-nucleating peptides VNp6 and VNp15 (a truncated VNp6); alongside a no-signal control. Figure 3B shows the six signals that gave measurable product plus the no-signal (NS) reference; VNp6 gave no detectable product. Each was fused upstream of the SUMO-peptide construct. The screen used four Target 6-binding designs (32 constructs), with 6 ml cultures, expressed under auto-induction for 16 h, at 25 °C.

**Figure 3.**
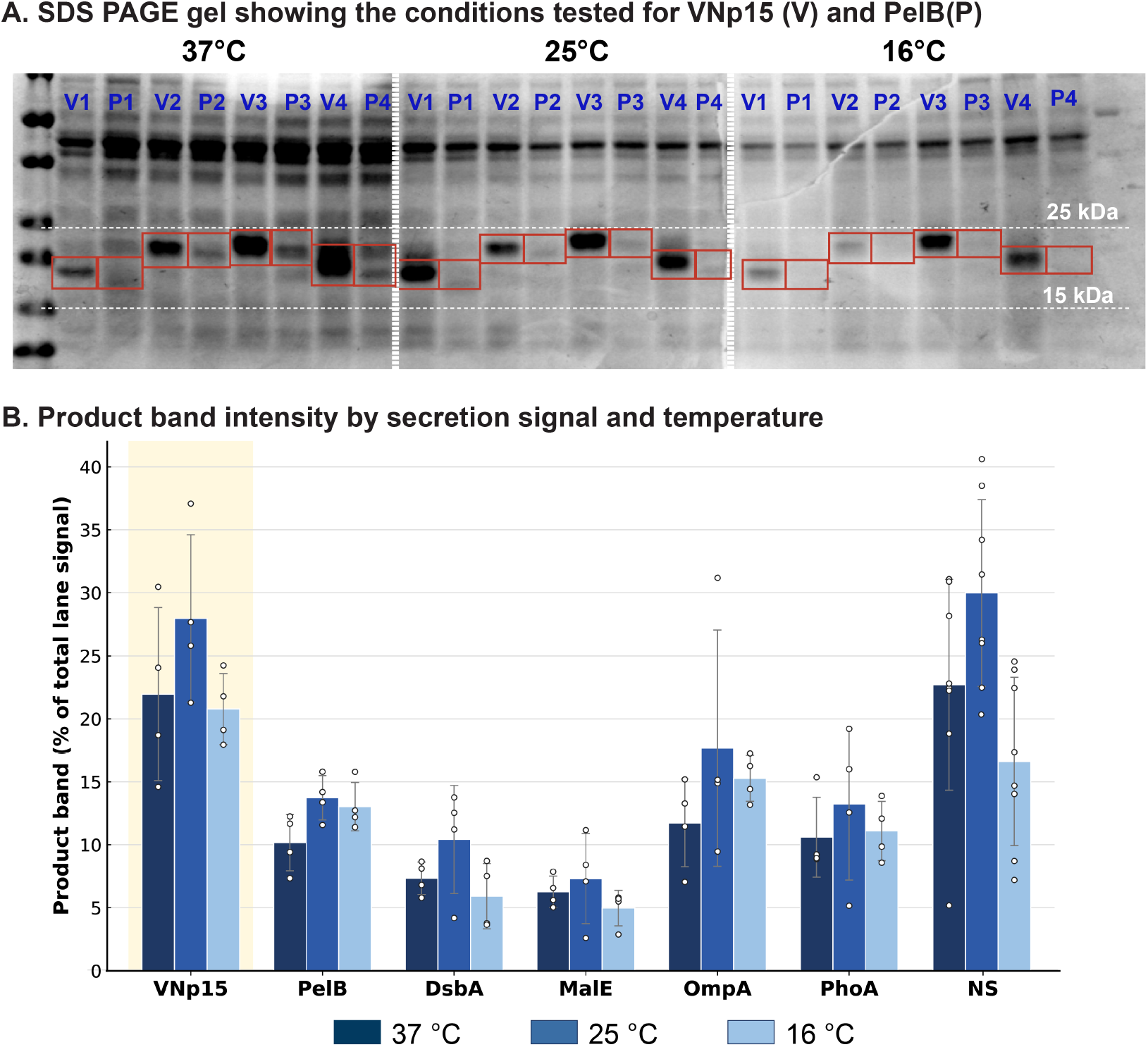
Secretion-signal and temperature optimisation of recombinant peptide export. **(A) Coomassie-stained SDS-PAGE** comparing the VNp15 (V) and PelB (P) secretion signals at three induction temperatures (37, 25 and 16 °C). Four representative designs were tested per signal, loaded in alternating lanes within each block (V1–V4, P1–P4). The white box marks the host reference band; red boxes mark the recombinant product near the 15 kDa marker (His6–SUMO–peptide). Markers at 25 and 15 kDa are indicated; product identity was confirmed by mass spectrometry. **(B) Densitometry** of the product band across all seven conditions and three temperatures. For every lane the boxed product band was integrated and expressed as a percentage of the total signal in that lane, so each value is internally loading-normalised. Bars show the mean of four designs (± 1 SD; individual designs overlaid, n = 4). Among the secretion signals tested, VNp15 gave the highest product fraction at every temperature and peaked at 25 °C. NS is a whole-cell extract of the untagged peptide, shown as a band-position reference (not a periplasmic fraction).

To extract the periplasmic content, we compared two periplasmic extraction methods compatible with 96-well throughput: *(i)* sodium deoxycholate (DOC) extraction (0.75% w/v DOC; 4 h at room temperature)^16^; and *(ii)* osmotic shock (a sucrose-based hypertonic-to-hypotonic transition)^17^. Cells were harvested and exposed to the lysis solutions as per protocols cited. Fractions of periplasmic content and remaining cell content were collected and resolved by *SDS-PAGE* and verified by *anti-His* Western blots. We observed a higher return of periplasmic content from the DOC protocol across the signal panel, and it required a single buffer exchange with less hands-on time. Therefore, we adopted the DOC protocol for routine use thereafter. Of the signals tested, VNp15 gave the highest periplasmic accumulation and was the only signal to give a clearly contrasting band and identity was confirmed by anti-His western blot against a positive control (Supplementary Figure 1).

As VNp15 gave the highest periplasmic yield, it was used for all subsequent expression optimisation trials. Four representative peptide designs were expressed with each of the six signals and compared across three induction conditions, with a no-signal construct as a baseline (Fig. 3A, B). 6 ml cultures were grown and expressed under auto-induction at three temperatures and time frames (16 °C for 16h, 25 °C for 16h, 37 °C for 4h). Expression was signal-dependent, with VNp15 returning the higher yields as percentage of total protein fraction (Fig. 3; Supplementary Figure 2). We observed higher yields and lower variation at 25°C for 16h and thereafter VNp15 was the chosen secretion signal for the system, with expression conditions following auto-induction at 25 °C for ~16 h, and DOC lysis for 4 h at room temperature.

### 4. Library Purification and Yield Analysis

To establish the pipeline at protocol-development scale, we adapted protocols developed by Qian, Milles, Wicky and colleagues^15^ based on IMAC and SEC onto our pipeline. We iterated this process until optimising it to the method described in Methods, section 2.4. We verified this method on a library of 60 disulphide-stapled peptide designs across nine protein targets (T1–T9), with 4-10 designs per target. All 60 designs gave purified peptide-fusion product above baseline.

Stacked SEC chromatograms showed a single dominant peak at a retention volume consistent with the expected ~17 kDa of the SUMO-tagged construct (Fig. 4A). Hierarchical clustering of normalised traces grouped designs by peak shape and showed that most profiles were similar, with a smaller subset showing earlier-eluting or shoulder-bearing peaks indicative of dimeric or aggregated species. The soluble-yield distribution (Fig. 4B) was strongly bimodal on a log scale, with a high-expression cluster around 150 μg per 6-mL culture and a low-expression cluster below 10 μg, and a library median of 135 μg post-pooling. The two well-separated populations indicate that the pipeline distinguishes productive from unproductive designs and provides a quantitative read-out for prioritisation.

**Figure 4.**
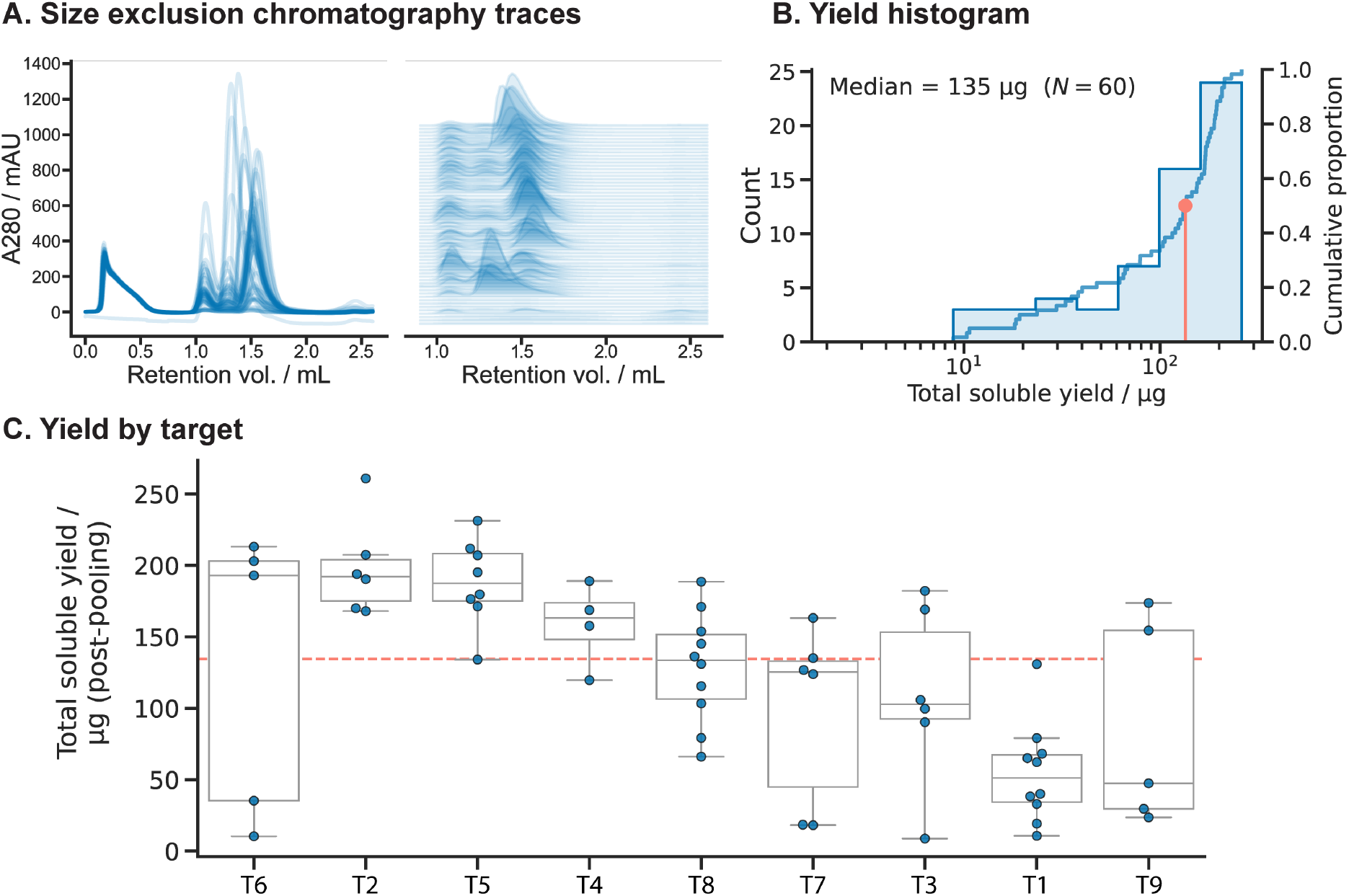
Library-scale purification and yield across a 60-peptide, nine-target panel. Sixty designed peptides spanning nine protein targets were carried through the fixed protocol: periplasmic secretion, IMAC, then size-exclusion chromatography (SEC) with no other deviation. **(A) SEC traces** (absorbance at 280 nm versus retention volume; Superdex 75 5/150, TBS) for all 60 peptides. ***Left***, traces overlaid on a common axis; ***right***, stacked traces with a vertical offset visualises individual profiles, showing that most peptides resolve as a single dominant peak in the expected elution window. **(B) Distribution of total soluble yield**: histogram (counts, left axis) and cumulative distribution overlaid (right axis). Yields range from 9 to 261 μg, median 135 μg (red marker); the x-axis is log-scaled to span the range. **(C) Total soluble yield after SEC pooling**, grouped by target (T; μg). Points are individual peptides; boxes show the median and interquartile range and whiskers extend to 1.5× the interquartile range. Targets are ordered by median yield (overall median 135 μg, range 9– 261 μg).

Yield varied by target (Fig. 4C). Designs against target 6 and 2 reached the highest median soluble yield (~190 μg), with those against T5 (~187 μg) and T4 (~163 μg) also strong. Designs for targets T8, T7 and T3 gave intermediate yields (~100–135 μg), and T1 and T9 the lowest (~50 μg). Across the library, 98% of designs yielded ≥10 μg, 78% ≥50 μg, 65% ≥100 μg and 45% ≥150 μg of purified peptide, ample for the microgram-scale binding assays used here. The per-target differences likely reflect intrinsic sequence properties (charge distribution, hydrophobicity, cysteine spacing) and match the target-dependent expression seen across protein-design pipelines^18^.

Plotting normalised SEC retention against expected molecular weight confirmed monomeric behaviour for most designs (Supplementary Figure 3), with a small fraction at retention values consistent with dimers or higher oligomers. Intact-mass analysis by mass spectrometry confirmed the expected mass of the His6– SUMO–peptide fusion within ±1 Da, confirming correct sequence identity. The protocol was later applied at scale to a separate 96-member panel by a different researcher, yielding soluble product for 92 of 96 designs (96%; median 95 μg; Supplementary Figure 5) and showing that it generalises beyond the development library reported here.

**Figure 5.**
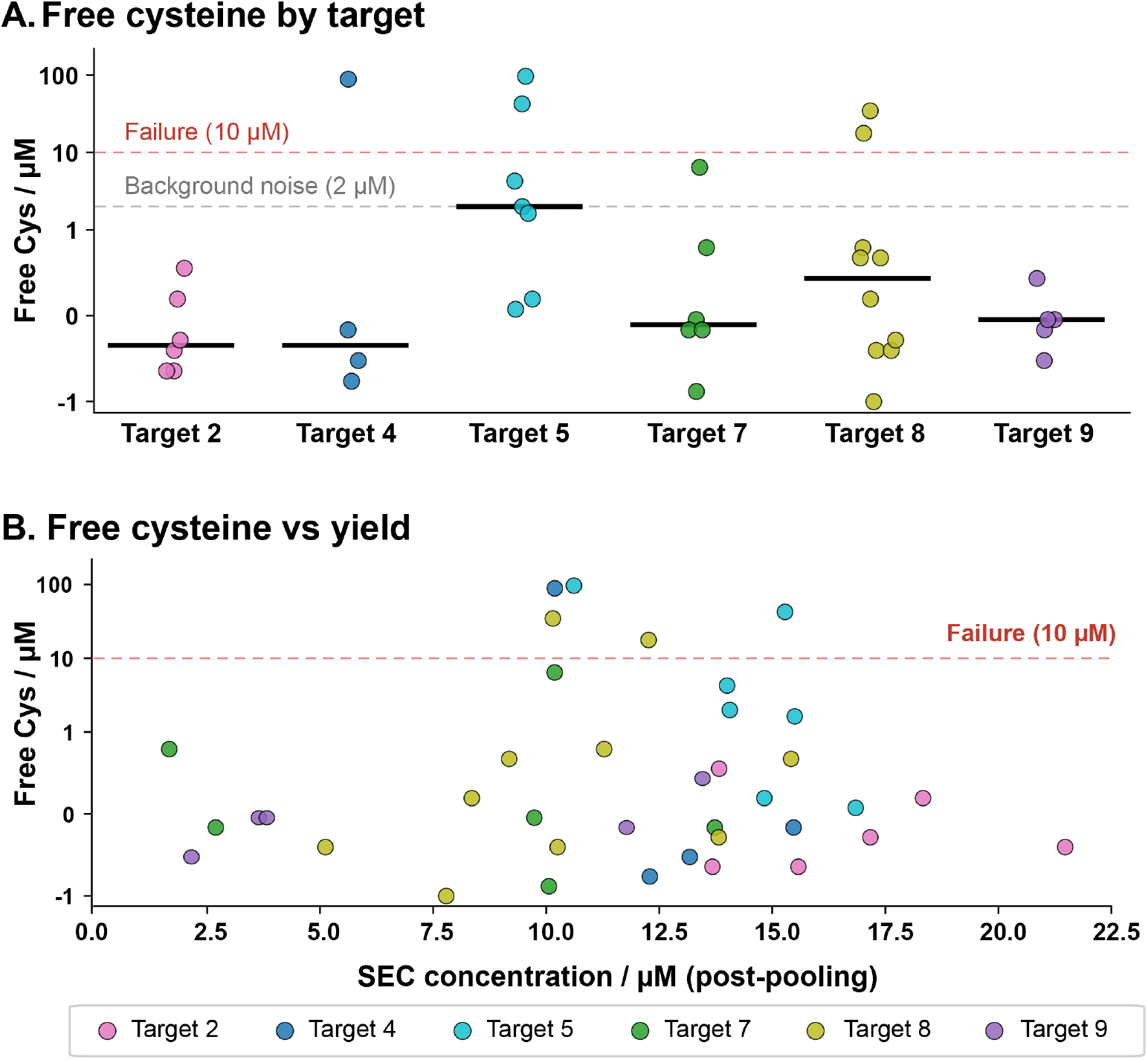
Disulphide staple formation assessed by Ellman’s assay. Free cysteine was measured for each purified peptide with Ellman’s reagent (DTNB); a low value indicates a formed disulphide staple, a high value an unformed one. **(A)** Free cysteine per peptide, grouped by target (n = 38 across six targets). Each point is one peptide; the black bar is the per-target median. Dashed lines mark the background-noise floor (2 μM) and the failure threshold (10 μM). Most peptides fall at or below the noise floor, indicating complete staple formation, with failures confined to a few designs. **(B)** Free cysteine against recovered concentration after SEC pooling, for the same peptides, coloured by target. Free cysteine does not track with yield: staple failure reflects the peptide sequence, not low expression.

### 5. Disulphide staple formation readout by Ellman’s assay

To assess intramolecular disulphide formation across the library, we ran Ellman’s assay^19, 20^ on a subset of 38 designs spanning six targets (T2, T4-5, and T7–9), against a freshly prepared 8-point L-cysteine standard curve (1–50 μM, R^2^ = 0.998 in the linear range; Supplementary Figure 4).

Free-thiol concentration in the well was read by interpolation onto the standard curve and compared with the peptide concentration from the SEC integral (Fig. 5A, B). Most designs (31/38; 82%) showed free-thiol concentrations below 2 μM within the assay baseline, consistent with full intramolecular disulphide formation. 5 of 38 designs (13%) gave free cysteine above 10 μM, indicating failed staple formation (T4: 1/4, T5: 2/7, T8: 2/10; T9, T2 and T7: 0). Per-target failure ranged from 0% (T9, T2, and T7) to 20% (T8), 25% (T4) and 29% (T5). Free-thiol concentration was uncorrelated with expression yield: the poorly stapled peptides were not necessarily those showing poor expression (Fig. 5B). Staple formation therefore depends on the sequence, not on expression level. The Ellman’s assay reports whether the disulphide staple formed, and whether a peptide that purifies remained unstapled. Designs with unformed disulphides can be deprioritised before binding assays, reducing instrument time spent on false negatives.

### 6. Recombinant pipeline produces target-binding peptides

To confirm that pipeline-produced peptides retain target-binding activity, representative recombinant (SUMO-tagged) constructs were compared with their corresponding peptide-only forms by surface plasmon resonance (Figure 6). A representative Target 5 design, T5-D1, bound in both the recombinant SUMO-tagged and peptide-only formats, with fitted *K*D values of 16.3 μM and 55 μM, respectively. In contrast, the corresponding linear peptide showed no detectable binding, supporting the relevance of the conformational constraint for target recognition.

**Figure 6.**
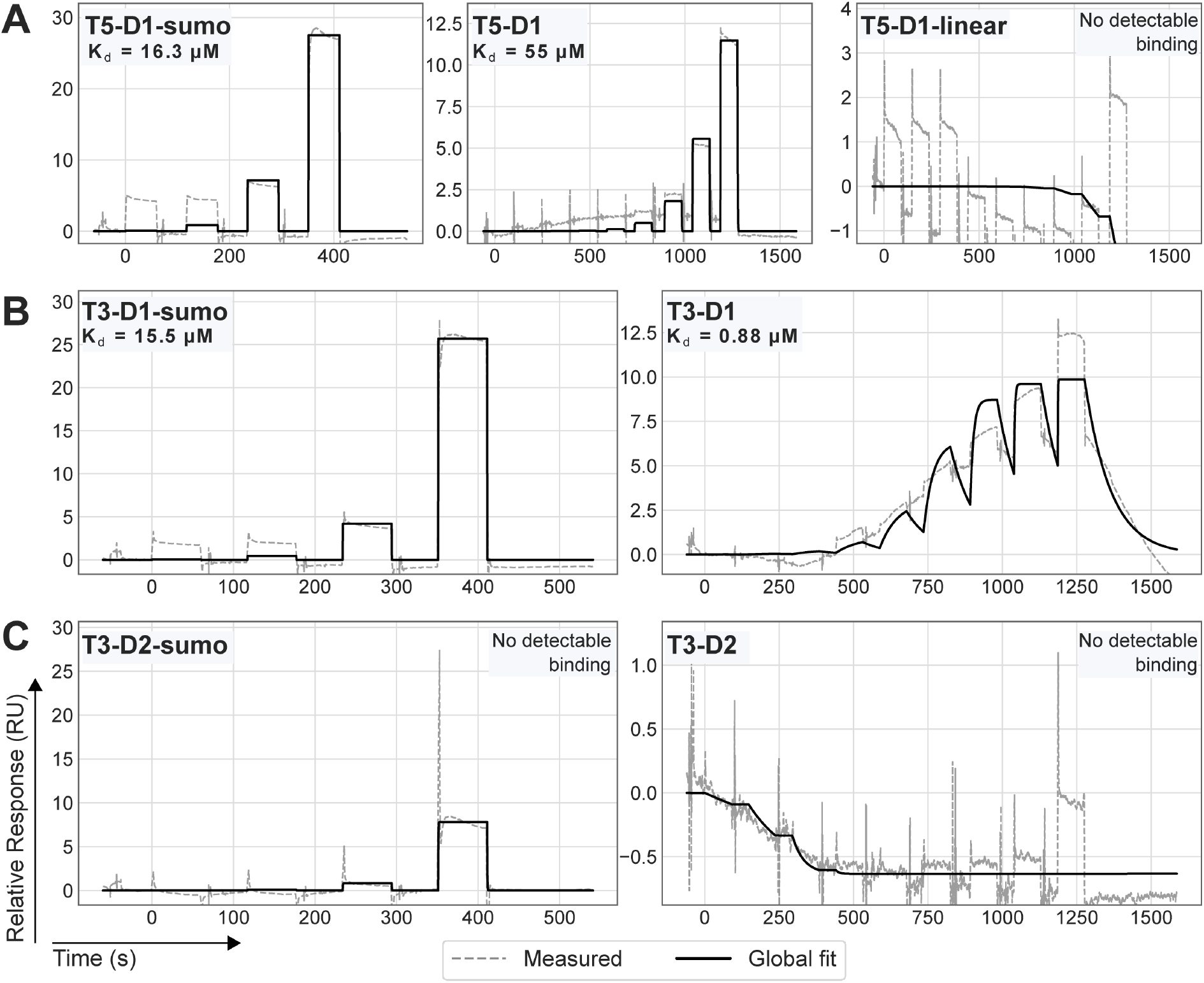
Matched SPR comparison of recombinant SUMO-tagged and peptide-only peptide formats. Surface plasmon resonance sensorgrams for representative peptide designs tested against their immobilised targets. Coloured traces show measured response and black traces show the fitted response where binding was detected. **(A)** Target 5 design T5-D1 tested in three molecular formats: recombinant SUMO-tagged construct (*K*_D_ = 16.3 μM), peptide-only form (*K*_D_ = 55 μM), and the corresponding linear peptide, which showed no detectable binding. **(B)** Target 3 design T3-D1 bound in both recombinant SUMO-tagged and peptide-only formats, with fitted *K*_D_ values of 15.5 μM and 0.88 μM, respectively. **(C)** Target 3 design T3-D2 showed no detectable binding in either SUMO-tagged or peptide-only format.

A second representative design, T3-D1, also bound in both formats, with a fitted *K*_D_ of 15.5 μM for the recombinant SUMO-tagged construct and 0.88 μM for the peptide-only form. This indicates that target binding is retained after recombinant production, although the apparent affinity can differ between the SUMO-tagged and peptide-only formats. As a matched negative control, a second Target 3 design, T3-D2, showed no detectable binding in either the SUMO-tagged or peptide-only format. These data show that the recombinant pipeline yields material suitable for direct binding screening and that SUMO-tagged constructs can preserve the binding behaviour of their corresponding peptide designs, while revealing design-dependent differences in apparent affinity.

## DISCUSSION

The pipeline delivers purified, quality-controlled disulphide-stapled peptide material for peptide design campaigns across a diverse set of targets. End-to-end execution from primer ordering to characterised material takes ~1 week of mostly hands-off plate work, at a per-variant DNA cost of ~$2. Applied to a 60-design library across nine targets (T1–T9), it gave soluble, monomer-shaped product for ~98% of designs above 10 μg and 65% above 100 μg, with a median total soluble yield of ~135 μg per 6-mL culture (range ~9–261 μg), sufficient for the microgram-scale binding assays reported here. Applied independently to a separate 96-member panel of optimised binders against one target, the pipeline reproduced this success (92 of 96 designs; Supplementary Figure 5), showing that it generalises beyond the development library.

The pipeline was adapted from the SAPP/DMX platform of Qian, Milles, Wicky and colleagues^15^, which addresses an overlapping but distinct problem. SAPP targets general recombinant production of *de novo*-designed proteins at hundreds of designs per day, runs end-to-end in 48 h, and with DMX oligo-pool demultiplexing, reaches ~$5 per construct using gene fragments at ~$0.07/bp. It reports 89.3% clonal purity across 929 GGA reactions and the expected mass within ±1 Da for 90% of 863 products and frames itself as a screening tool. Our pipeline is built for a class of biomolecule that SAPP does not extend to: cyclic disulphide-stapled peptides, whose production needs translocation to an oxidising compartment, cysteine pairing during or after folding, and a QC read-out for the staple; none of which yield, dispersity or intact mass report. The workflows are complementary. The reliability figures fall in the same range: ~82% of measured designs passed the Ellman’s threshold here and ~90% of wells expressed above 2 μM, close to SAPP’s 90% mass-correct rate, and consistent with incomplete well-level success being the expected behaviour of any screening-scale recombinant pipeline.

Multi-target validation is a strength of the dataset. Because the 60-design library spans nine targets with 4–10 designs each, per-target yield distributions can be compared directly: T2, T5, T6 and T4 cluster at the high end (medians ~160–190 μg), T1 and T9 at the low end (medians ~50 μg), with the other targets between. This spread indicates that the rate-limiting step for yield is sequence-and target-specific, likely hydrophobicity, predicted aggregation propensity, or translocation efficiency^18, 21^, and not particular to any one target. The workflow performs across all nine targets without per-target re-optimisation, as required of a generalisable methods pipeline.

Among the QC steps, Ellman’s detection of free sulfhydryl groups distinguishes this workflow from general-purpose recombinant pipelines. We measured free Cys for 38 of the 60 designs, across six of the nine targets (Targets 2, 4, 5, 7-9,). Failure to form the intramolecular disulphide ranged from 0% (Targets 2, 7 and 9) to 25–29% (Target 4 and 5), and free thiol was uncorrelated with SEC yield. A design can express well, run as a monomer by SEC, give the correct intact mass, and still fail to form the staple. Ellman’s detects staple failure that yield, dispersity and intact mass would miss; it therefore constitutes a distinct QC step for any disulphide-constrained peptide. The 38/60 coverage is a representative subset, spanning enough target classes to support the point that staple-formation failure is sequence-driven and decoupled from soluble expression.

The expressed peptides can be readily used in binding assays: surface plasmon resonance showed that the recombinant designs bind their targets (Figure 6). Matched comparisons with the corresponding peptide-only forms showed that binding activity can be retained in the SUMO-tagged format, although apparent affinity can differ between formats in a design-dependent manner. A matched non-binding design remained non-binding in both formats, while the corresponding linear form of one constrained binder showed no detectable binding, supporting the functional importance of the conformational constraint. Further, the intact ~17 kDa His6-SUMO fusion may be advantageous in label-free, mass-sensitive biosensor-based analysis, such as surface plasmon resonance and bio-layer interferometry. In these methods the response scales with the molecular weight of the analyte. A free stapled peptide is small, so at a given molar occupancy it returns a relatively low signal, and weak binding can be difficult to distinguish from the baseline. The His6-SUMO fusion is roughly 8-fold heavier than the free peptide (~17 kDa versus ~2 kDa for a 16–25-residue design); the same binding event is therefore expected to generate a proportionally larger response, which can facilitate measurement of weaker interactions.

Several limitations should be stated. Functional binding was examined in a limited set of matched SUMO-tagged and peptide-only designs; these comparisons showed that binding activity can be retained but also revealed design-dependent differences in apparent affinity, so a systematic survey across more targets would be needed to determine how generally the SUMO fusion preserves quantitative peptide-only affinity. Oxidative folding in the periplasm is not guaranteed for all topologies; designs needing non-disulphide constraints, non-canonical residues, or multiple disjoint disulphide bonds fall outside the present scope. Finally, the workflow has been validated to ~96 samples in a single plate-format run, and scaling beyond a few plates per week will likely need liquid-handling automation similar to DMX^15^.

Coupling to active-learning design loops^22^ would connect computational generation with experimental read-out, with Ellman’s pass/fail and SEC yield as immediate sequence-level labels for retraining. Adapting the workflow to other disulphide-constrained formats, such as, bicyclic and multicyclic topologies, or cystine-knot (knottin) miniproteins, would widen the chemical scope while reusing the cloning, expression and SEC modules unchanged. Broader target validation, especially against intracellular protein–protein interfaces and shallow allosteric pockets, would test the limits of the design–express–QC loop. Together these point toward sequence-encoded macrocyclic peptide libraries designed, expressed, quality controlled and screened on a timescale matched to modern generative design.

## MATERIALS AND METHODS

### 1. *In vivo* Assembly (IVA) Cloning

Peptides were cloned into plasmids using *in vivo* assembly (IVA) cloning^14^. Following this method, peptide sequences are introduced via a PCR reaction with primers that anneal to extend a sequence that incorporate a peptide-encoding sequence and deletes any sequences that are not amplified. The assembly methods were optimised for production in medium throughput, enabling parallel processing of 96 samples, following the medium-throughput scaling approach of Qian et al.^15^.

#### 1.1. Construct Design

A synthetic DNA insert was designed to include a bacterial secretion signal, a GWGS linker, a hexahistidine (6×His) tag, an SGGDR linker, a Small Ubiquitin-like Modifier (SUMO) domain, a WGGSSGG–Cys linker, the peptide-insertion site, a C-terminal cysteine, and a stop codon (the control-of-cell-death B (ccdB) selection gene occupies the peptide-insertion site in the parent vector and is replaced by the peptide during IVA cloning). The two cysteines that form the disulphide staple are encoded by the construct, flanking the peptide-insertion site; one in the fixed WGGSSGG–Cys linker downstream of SUMO and one introduced by the primer at the peptide C-terminus. The designed peptide is inserted between them, so the staple forms across the variable sequence between two scaffold-encoded cysteines (see Fig. 2). The cysteines that serve as staples are included in the construct. The construct was cloned into a pET24b(+) vector, featuring a T7 promoter under LacO regulation, a high-efficiency ribosome binding site, a T7 terminator, and a kanamycin resistance cassette.

#### 1.2. Primer Design

Primers were designed in three regions: (i) a fixed 3′ adapter tail, (ii) a peptide-specific extension region, and (iii) a 5′ homology block (see Fig. 2A). The 3′ adapter tail anneals to the plasmid template, the extension region introduced the peptide-encoding sequence during amplification, and the 5′ homology block enables the circularisation of the amplified plasmid by triggering RecA-independent homologous recombination. The 3’ adapter tails were designed to anneal upstream and downstream of the *ccdB* gene, to facilitate its removal during amplification. This ensures that only bacterial cells transformed with the amplified vector, carrying the peptide sequence instead of ccdB, survive the selection.

The forward primer adapter tail (5′-TGTTGACGCTGAGCCGGAGT-3′) included a Stop Codon (TGA). The reverse primer adapter tail (5′-ACAACCTCCCGAGCTGCCT-3′) annealed to the region encoding the Cys– Gly_2_–Ser_2_–Gly (CGGSSG) residues of the **WGGSSGGCG** linker. Both primers incorporated cysteine codons facilitating the disulphide bridges and both adapter tails were designed with a Tm of 63 °C to enable a two-step PCR.

A Python-based pipeline was developed to automate the design of primers for 96-well peptide libraries (see extended data). Amino acid sequences were reverse-translated and codon-optimised for E. coli using the E. coli standard genetic code via Biopython. DNA sequences were then optimised with DNA Chisel to satisfy the following constraints: (i) a global GC content of ~0.60; (ii) avoidance and removal of homopolymers (runs ≥ 5 nts) and long GC/AT-rich segments (> 8 nts); (iii) removal of restriction sites that could interfere with downstream processes; and (iv) avoidance of secondary structures at the mRNA level that could affect translation initiation.

The workflow automatically generates forward and reverse primer pairs consisting of three distinct functional regions: (i) a fixed 3′ adapter tail with a melting temperature of 63 °C for template annealing; (ii) a central peptide-encoding extension; and (iii) a 5′ homology block. The length of the 5′ homology block was iteratively calculated (ranging from 14 to 20 nts) to target a melting temperature (Tm) between 48 °C and 56 °C, ensuring efficient RecA-independent homologous recombination during the IVA process (see figure 2).

The workflow yields csv files mapping each sequence to a specific coordinate in a 96 well plate and calculates the theoretical molecular weights and lengths for each expressed peptide fusion to provide a reference for the subsequent analyses. Further, the files can be directly used to order from a provider. In this study primers were acquired from Integrated DNA Technologies (Coralville, Iowa, USA) as 500 picomoles of custom DNA oligos.

#### 1.3. Real Time Polymerase Chain Reaction (qPCR) Amplification

DNA oligonucleotides were eluted in nuclease-free ultrapure water to 5 μM. Each 10 μL qPCR reaction contained 5 μL of 2× Q5 Master Mix (NEB, Cat. No. M0492), 0.5 μM of each assembly primer, plasmid template at 0.8 ng/μL (8 ng per reaction), 1× EvaGreen (Biotium, 31000) and nuclease-free water to volume, following the Benchling protocol (“Cloning of ds-stapled peptides for expression in E. coli”). Reactions were prepared on ice in 96-well low-profile unskirted white qPCR plates (Bio-Rad, Cat. No. MLL9651). The forward and reverse primers were dispensed first at the bottom of the plate, followed by the Master Mix, containing the plasmid template. The mixture was gently pipetted once to ensure homogeneity.

Amplification was performed using a CFX96 Real-Time PCR Detection System (Bio-Rad, Cat. No. 1855095). The programme included an initial denaturation at 98 °C for 30 s, followed by 20 cycles of 98 °C for 10 s and combined annealing/extension at 72 °C for 140 s, with a final extension at 72 °C for 2 min. Yields were considered sufficient for transformation if the exponential amplification phase was reached before cycle 15 and exceeded 5,000 relative fluorescent units (RFU).

#### 1.4. Removal of template plasmids

Parental methylated DNA was digested using DpnI (New England Biolabs, Cat. No. R0176). The enzyme was diluted in EB buffer (10 mM Tris-HCl, pH 8.0) to 4 U/μL and added to the PCR products. The reaction was incubated at 37 °C for 45 min, followed by heat inactivation at 80 °C for 20 min.

#### 1.5. Transformation

For each 96-well plate of PCR products, eight 200 μL aliquots of chemically competent E. coli BL21(DE3) (New England Biolabs, Cat. No. C2527I) were thawed on ice for ~15 min. In parallel, 1 μL of each PCR product was dispensed into a pre-chilled Applied Biosystems MicroAmp™ plate (Thermo Fisher Scientific, Cat. No. N8010560) held on a cooling rack, using a 0.5–10 μL pipette.

Each cell aliquot was gently resuspended by tapping the tube lightly 3–4 times. Using pre-chilled 2–20 μL pipette tips (stored at −20 °C), 15 μL of competent cells were added directly into the DNA droplets and mixed gently by swirling the tip at the bottom of the well 2-3 times. Throughout setup, all components were kept on ice (4 °C). The plate was incubated on ice for exactly 30 min without disturbance. Heat shock was performed for 20 s at 42 °C by placing the sealed plate in a calibrated water bath with uniform well submersion. The plate was immediately returned to ice for 5 min.

For outgrowth, 100 μL SOC medium (New England Biolabs, Cat. No. B9020) at room temperature was added to each well, the plate was sealed with a sterile breathable film, and cells were recovered for exactly 1 h at 37 °C shaking in a plate incubator (with continuous agitation). For selection, 100 μL of the recovered mixture was transferred into a deep-well plate containing 900 μL of LB supplemented with kanamycin (50 μg/mL). Plates were sealed with breathable film and incubated at 37 °C with vigorous shaking overnight (≤16 h).

### 2. Protein expression and purification

#### 2.1. Media and Buffers

Autoinduction medium was prepared using Terrific Broth (TB; 40 g/L) (MP Biomedicals, Cat. No. 113046032) and supplemented with 50 mM Na_2_HPO_4_, 50 mM KH_2_PO_4_, 100 mM NH_4_Cl, 10 mM Na_2_SO_4_, 0.05% (w/v) glucose, and 0.2% (w/v) α-lactose, as well as 2 mM MgSO_4_, 0.2× Trace Metals solution (Teknova, Cat. No. T1001), 10 μM thiamine, and kanamycin (100 μg/mL final). The final volume was adjusted to 1 L with ultrapure water, and the medium was sterilised by filtration, as previously described. Lysis buffer consisted of 100 mM Tris-HCl (pH 8.0) containing 0.75% (w/v) sodium deoxycholate (0.563 g DOC per 75 mL of buffer) and 1× cOmplete™ EDTA-free Protease Inhibitor Cocktail (Roche, Cat. No. 11836170001). The buffer was mixed until fully dissolved and kept on ice until use. Wash buffer contained 20 mM Tris-HCl (pH 8.0), 250 mM NaCl, and 10 mM imidazole. Elution buffer contained 20 mM Tris-HCl (pH 8.0), 500 mM NaCl, and 500 mM imidazole.

#### 2.2. Expression

For expression, 100 μL of the overnight culture was used to inoculate 900 μL of auto-induction media with kanamycin in a fresh 96-deep-well plate. In a typical experimental batch, 6–8 plates were processed simultaneously. The plate was sealed with a gas-permeable membrane and incubated at 25 °C for ~16 h shaking at 300 rpm.

#### 2.3. Lysis

Cells were harvested by centrifugation at 2,000 × g for 5 min at 8–10 °C to minimise cell wall disruption. The supernatant was removed by quick inversion and blotting on absorbent paper. To manage the volume for periplasmic content extraction, the library was consolidated through sequential washes and pooling. Cell pellets from the original expression plates were washed once with 400 μL of 100 mM Tris-HCl, pH 8.0 supplemented with 100 mM N-ethylmaleimide (NEM) to trap free thiols during processing and cOmplete EDTA-free protease inhibitor cocktail (Sigma-Aldrich), mixed gently by swirling or light vortex until fully resuspended and the library was pooled into four plates. This was repeated once more to pool the library onto two plates.

Lysis followed the sodium deoxycholate periplasmic-extraction method^16^. The consolidated pellets were resuspended in 350 μL of lysis buffer and incubated for 4 h at room temperature on an orbital or lateral shaker at low speed (approximately 35 rpm). Lysates were clarified by centrifugation at 4,000 × g for 25 min at 8 °C, and 200–250 μL of the clear supernatant was carefully collected (without disturbing the pellet) and transferred to one clean plate.

#### 2.4. Immobilised Metal Affinity Chromatography (IMAC)

For each 96-well plate of clarified periplasmic extract, 10 mL of 50% (v/v) Ni–NTA resin slurry in 20% (v/v) ethanol was equilibrated on a 96-well polypropylene fritted plate (800 μL/well, 25 μm; Agilent Technologies, Cat. No. 200953-100) by sequential washing with 20 column volumes (CV) of ultrapure water followed by 20 CV of equilibration buffer (20 mM Tris-HCl, pH 8.0, 250 mM NaCl, 10 mM imidazole). Buffer removal between steps was performed using a vacuum manifold.

Using a multichannel pipette, 150 μL of the equilibrated 50% resin slurry (corresponding to ~75 μL settled resin; binding capacity ~0.75 mg His-tagged protein) was dispensed into each well containing clarified periplasmic extract. The resin–lysate mixtures were transferred to a clean 96-well conical-bottom polypropylene plate (Nunc® MicroWell™; Merck, Cat. No. P7241) and incubated for 40 min at 4 °C on a slow orbital or lateral shaker to allow binding. The mixtures were then transferred back onto the fritted plate and the flow-through was removed under vacuum.

The resin was washed 4–5 times with 800 μL wash buffer, applying vacuum after each wash and avoiding over-drying of the resin bed. The residual wash buffer was removed by briefly centrifuging the fritted plate at 500 × g for 5–10 s. Bound proteins were eluted by adding 200 μL of the elution buffer and incubating for 2–3 min. Eluates were collected by centrifugation at 500 × g for 2 min into a clean 400 μL flat-bottom receiver plate. Eluates were subsequently filtered using a 0.2 μm PTFE 96-well filter plate (Pall Life Sciences, AcroPrep Advance 2 mL, 0.2 μm PTFE membrane) placed on top of a conical-bottom 96-well polypropylene plate (Nunc® MicroWell™; Merck, Cat. No. P7241) and centrifuged at 4,000 × g for 10 min.

#### 2.5. Size Exclusion Chromatography (SEC)

Filtered eluates were analysed by size exclusion chromatography (SEC) using an Agilent HPLC system (Agilent 1260 Infinity II LC System). Separations were performed on a Superdex 75 5/150 column (~3 mL bed volume; Cytiva). Elution was monitored by UV absorbance at 280 nm to assess sample purity and estimate peptide yields. SEC chromatograms were processed with a custom Python analysis built on the SAPP/DMX framework, shared in extended data^15^ to align traces across runs and map each chromatogram to its corresponding 96-well library coordinate. As shown in figure 4, for the stacked-trace display, normalised absorbance profiles were grouped by average-linkage (UPGMA) hierarchical clustering with a Euclidean distance metric (fastcluster). Fractions were eluted in TBS.

### 3. Detection of Free Sulfhydryl Groups

Free sulfhydryl groups were quantified using Ellman’s assay^19, 20^. Reaction buffer (RB) consisted of 0.1 M Tris-HCl (pH 8.0). A DTNB stock solution (10 mM) was prepared by dissolving 40 mg of 5,5′-dithiobis(2-nitrobenzoic acid) (DTNB; Sigma-Aldrich, D8130; MW 396.35) in 10 mL DMSO. Immediately before use, the DTNB stock was diluted 1:200 in RB to obtain a 50 μM DTNB working solution.

A cysteine standard curve was prepared using L-cysteine hydrochloride monohydrate (Sigma-Aldrich, C7880; MW 175.6). A 10 mM cysteine stock solution was prepared by dissolving 17.56 mg in 10 mL RB and serially diluted in RB to generate an eight-point standard curve (0–50 μM).

For the assay, 50 μL of each standard or protein sample (diluted in RB) was dispensed into a 96-well UV-STAR microplate (Greiner, 675801). Standards were run in triplicate and samples in duplicate. DTNB working solution (50 μL) was added to all wells (final volume 100 μL; final DTNB 25 μM), and plates were incubated for 15 min at room temperature with orbital shaking. Absorbance was recorded using a spectral scan from 300 to 500 nm, and quantification was performed at 412 nm with pathlength correction enabled. Free thiol concentrations were calculated using the molar extinction coefficient of 2-nitro-5-thiobenzoate (TNB; ε412 = 13,600 M^−1^ cm^−1^)^20^ and/or by interpolation from the cysteine standard curve. This measurement was used to assess intramolecular disulphide staple formation by comparing the free thiol content of stapled peptides against reduced controls. A custom Python script automated the calculations and returned thiol concentrations for each sample.

### 4. Surface Plasmon Resonance

Binding measurements were performed on a Cytiva Biacore 8K in HBS-EP+ running buffer at 25 °C. Biotinylated target proteins were captured on the sensor surface using the Cytiva Biotin Capture Kit, and recombinant His6-SUMO constructs, corresponding peptide-only forms, and linear peptide controls where indicated were injected as analytes at 30 μL min^−1^. Measurements were performed using single-cycle kinetics, with analyte concentration ranges selected according to the individual design and experiment. Sensorgrams were double-referenced using reference-surface and buffer-blank subtraction and analysed in Biacore Insight using a 1:1 binding model where concentration-dependent binding was observed. Conditions were as described previously^4, 5^.

### 5. Mass spectrometry

Intact mass spectra were obtained by reverse-phase LC/MS on an Agilent G6230B TOF using an AdvanceBio RP-Desalting column with a fast gradient of 90:10 to 5:95 (A:B) over 2 min (A: water + 0.1% formic acid; B: acetonitrile + 0.1% formic acid) and deconvoluted with BioConfirm using a total-entropy algorithm^15^. Observed intact masses of the His6–SUMO–peptide fusions were compared with values calculated from the design sequences.

## DATA AND CODE AVAILABILITY

The analysis code (automated primer design, cloning quality control, size-exclusion chromatography and Ellman’s assay), the expression construct sequence (His6–SUMO–linker–ccdB in a pET-24b(+) backbone), and sequence-free source data for Figures 2–5 are openly available at Zenodo (https://doi.org/10.5281/zenodo.23035383). Designed peptide amino-acid sequences are withheld as intellectual property. Surface plasmon resonance data for the representative recombinant SUMO-tagged constructs, corresponding peptide-only forms, and linear peptide control are included. The size-exclusion chromatography analysis builds on the SAPP/DMX framework^15^.

## USE OF AI TOOLS

During preparation of this work, the authors used an AI assistant (Claude, Anthropic) to assist with the code generating and refining the data-analysis and figures, packaging the reproducibility deposit, and editing portions of the text. The tool was not used to generate experimental data. All AI-assisted outputs were checked, tested, and approved by the authors, who are fully responsible for the content.

## AUTHOR CONTRIBUTIONS

Author contributions are reported using the CRediT taxonomy. Conceptualisation: Y.F.B.; Methodology: Y.F.B., G.G.A., G.B.; Investigation: Y.F.B., G.G.A., J.J., S.R., X.L.; Software: Y.F.B., G.G.A.; Formal analysis: Y.F.B., G.G.A.; Writing – original draft: Y.F.B., G.G.A.; Writing – review and editing: all authors; Supervision: G.B., M.T., D.B.; Funding acquisition: Y.F.B., G.B., M.T., D.B.; Author initials: Y.F.B. (Yensi Flores Bueso), G.G.A. (Gizem Gökçe-Alpkılıç), J.J. (Jihun Jeung), S.R. (Stephen Rettie), X.L. (Xinting Li), M.T. (Mark Tangney), D.B. (David Baker), G.B. (Gaurav Bhardwaj).

## FUNDING AND ACKNOWLEDGEMENTS

This work was funded by grants to G.B. (National Institutes of Health award number 5R21AI178088-02; the National Institute of General Medical Sciences of the National Institutes of Health under award number R35GM163670; and startup funds from the University of Washington’s Department of Medicinal Chemistry, the UW Institute for Protein Design and the Audacious Project) and to D.B., the Howard Hughes Medical Institute, HHMI. Y.F.B. was supported by the European Union’s Horizon Europe research and innovation programme under the Marie Skłodowska-Curie Actions grant agreement No 101059124 (BacStar). Views and opinions expressed are however those of the author(s) only and do not necessarily reflect those of the European Union or the European Research Executive Agency (REA). Neither the European Union nor the granting authority can be held responsible for them. The authors thank Ljubica Mihaljevic, Marc Exposit, Mohammad Abedi, Adam Chazin-Gray, Florian Praetorius, Basile Wicky, Jason Qian, and Adam Broerman for technical assistance and helpful discussions.

