## Supplementary Figures 1-5 for "A scalable recombinant pipeline for disulphide-stapled peptides"

#### Supplementary Information

A scalable recombinant pipeline for disulphide-stapled peptides validated across diverse target classes

A

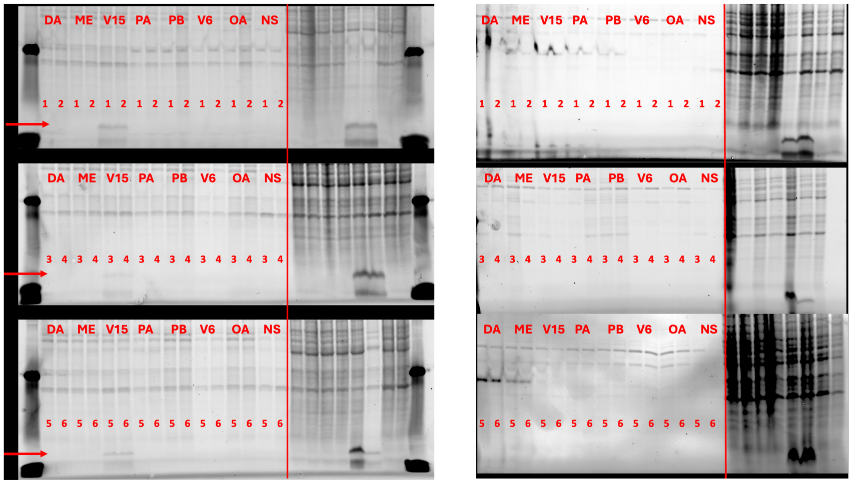


B

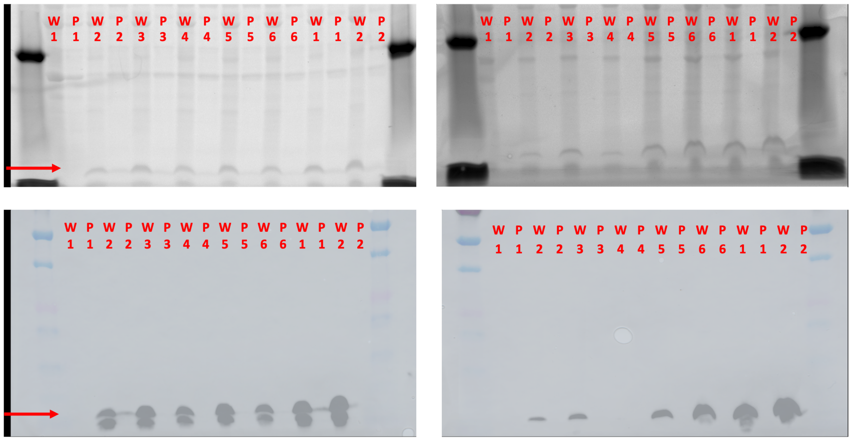


**Supplementary Figure 1. Periplasmic release across the signal-peptide panel and confirmation of product size.** **(A)** Stained SDS-PAGE comparing two periplasmic-release methods: sodium deoxycholate (DOC; left) and sucrose-based osmotic shock (right). Across the signal-peptide panel (lanes DA, ME, V15, PA, PB, V6, OA and the no-signal control NS). For each signal peptide the six peptide designs are numbered 1–6 across the three stacked gels; the red arrow marks the $His_6$–SUMO–peptide product band. (B) Confirmation of product identity and size. Paired whole-cell (W) and periplasmic (P) fractions for designs 1–6, followed by two untagged no-His controls, were resolved after DOC (left) and osmotic-shock (right) extraction. (Top) Coomassie-stained SDS-PAGE; (bottom) anti-$His_6$ western blot of the same lane layout, with the colour molecular-weight ladder overlaid, locating the $His_6$–SUMO–peptide product at ~15 kDa and confirming that the lower Coomassie band is the intended product.

#### Supplementary Information

A scalable recombinant pipeline for disulphide-stapled peptides validated across diverse target classes

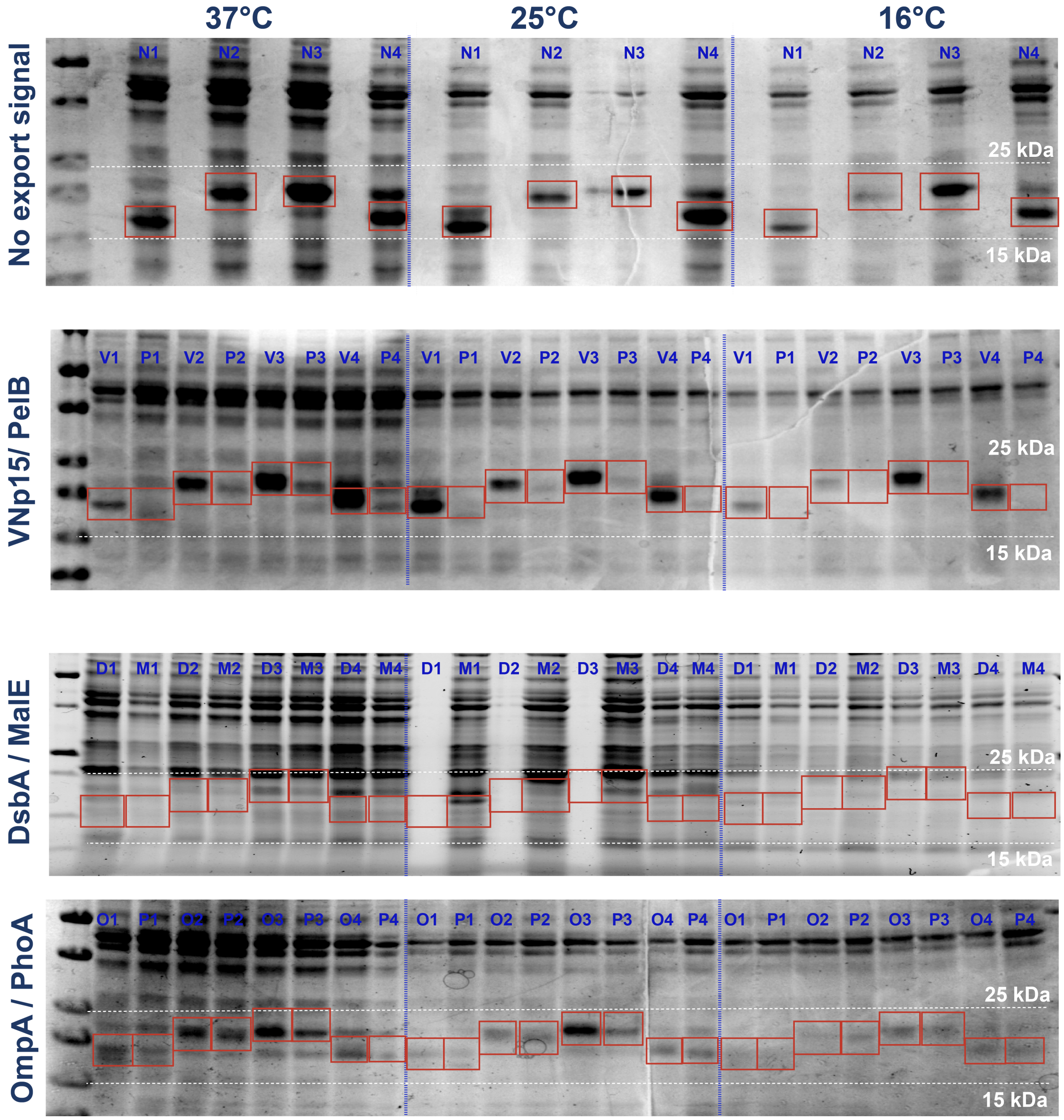


**Supplementary Figure 2. Secretion-signal screen across induction temperatures.** Coomassie-stained SDS-PAGE for the remaining secretion signals tested in the screen: DsbA and MalE, OmpA and PhoA, and a no-secretion signal (NS) control. Each at 37, 25 and 16 °C, completing the VNp15/PelB comparison shown in Figure 3. Boxed bands mark the recombinant product; the per-signal, per-temperature product fraction is quantified in Figure 3B.

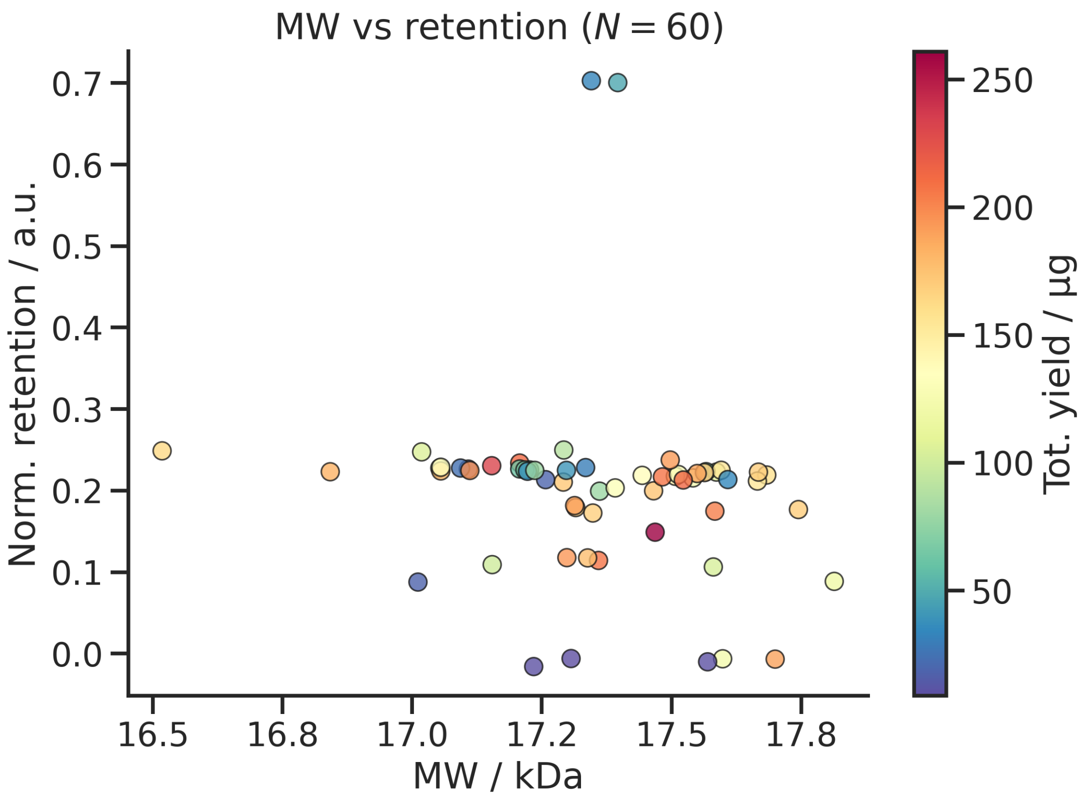


**Supplementary Figure 3. Size-exclusion retention versus expected molecular weight across the 60-design library.** Normalised SEC retention volume plotted against the calculated molecular weight of each $His_6$–SUMO–peptide construct. Most designs cluster at the retention expected for the ~17 kDa monomer, with a minority at earlier-eluting positions consistent with dimeric or higher-order species.

#### Supplementary Information

A scalable recombinant pipeline for disulphide-stapled peptides validated across diverse target classes

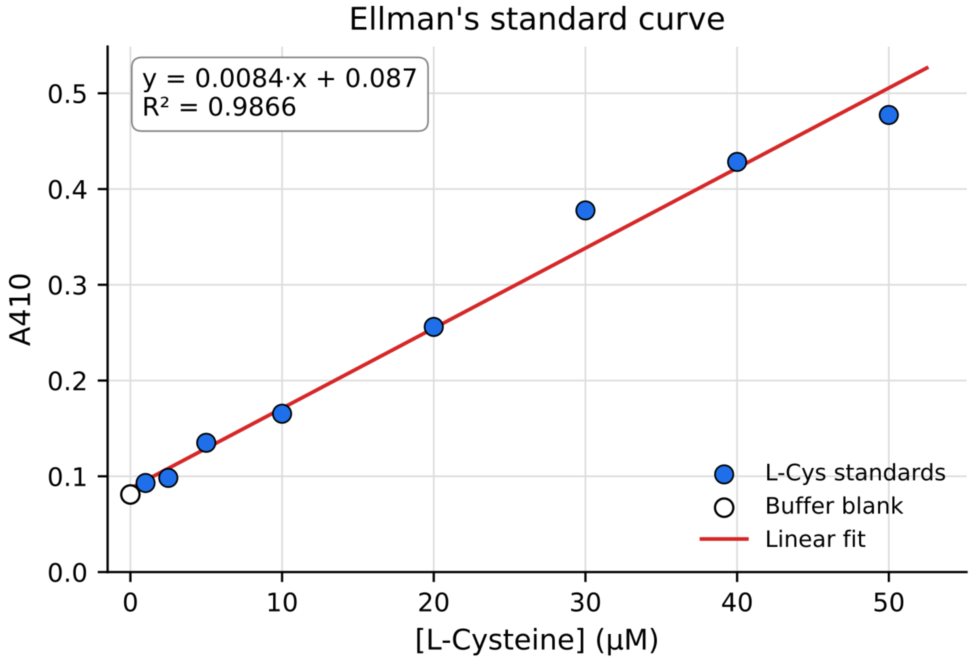


**Supplementary Figure 4. Ellman's assay L-cysteine standard curve.** Eight-point L-cysteine standard curve (1–50 µM) with linear fit ($R^2$ = 0.998 in the linear range), used to interpolate free-thiol concentrations for the disulphide-staple read-out (Figure 5).

### Supplementary Information

#### A scalable recombinant pipeline for disulphide-stapled peptides validated across diverse target classes

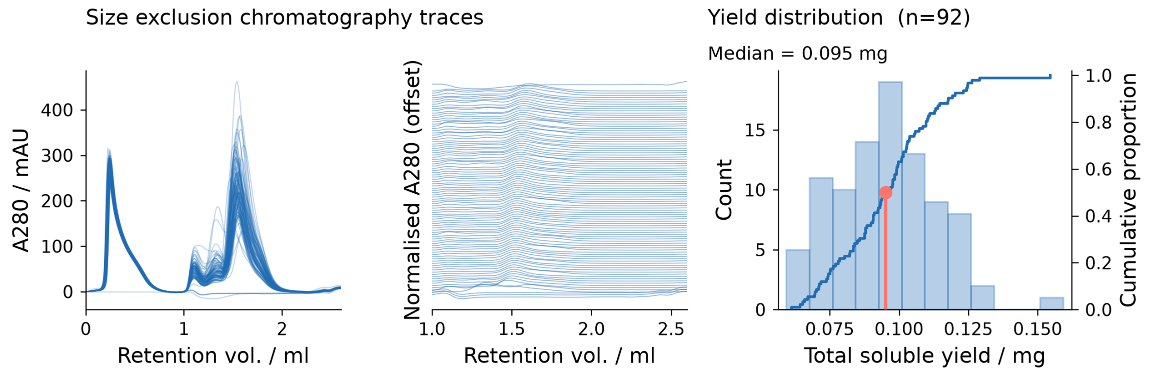


**Supplementary Figure 5. Purification and size-exclusion analysis of an independent, full-scale re-validation batch.** Site-saturation variants of an optimised binder against one the initial targets were produced through the pipeline in a single 96-well batch by an independent operator. (Left) Overlaid analytical size-exclusion chromatography traces (A280) for the recovered designs. (Middle) The same traces, baseline-offset and normalised, showing consistent, predominantly monomeric elution. (Right) Distribution of total soluble yield across the 92 of 96 designs that gave recoverable soluble product (median 0.095 mg). The run reproduces pipeline performance at full 96-well scale on a design set independent of the 60-design development library.
